# The extracellular matrix in selective decussation of retinal ganglion cell axons: β2 laminins regulate the ipsilateral projection

**DOI:** 10.64898/2025.12.28.696306

**Authors:** Alanis Hernandez-Arce, Madeline Turo, Adam N. Robinson, Skylyn McNamara, Danny Yeo, Reyna I. Martínez-De Luna

**Author notes:** Lead contact: Reyna I. Martínez-De Luna.

## Abstract

In binocular animals, retinal ganglion cell (RGC) axons selectively decussate at the optic chiasm. Selective decussation is directed by radial glia and midline neurons in the optic chiasm. Radial glia attach to the underlying pial basement membrane (PBM) that is rich in β2 laminins. Here, we asked whether β2 laminins in the PBM control the selective decussation of RGC axons. Genetic deletion of the β2 subunit increased the proportion of non-decussating RGC axons in the ipsilateral tract. β2 laminins are expressed in the PBM during the peak and late phases of RGC axon growth, and their deletion results in fragmentation of the PBM and dysmorphic radial glia. Consistent with the increase in the proportion of ipsilateral axons, we found persistent expression of the ipsilateral guidance cue EphrinB2 in radial glia during the late phase of axonal decussation, thus extending the developmental window for the development of the ipsilateral projection. Surprisingly, this increase in EphrinB2 was accompanied by an increase in the number of ipsilateral RGCs in the ventrotemporal retina. These results suggest that β2 laminins regulate the size of the ipsilateral projection by providing cues that control the generation of ipsilateral RGCs and the expression of the ipsilateral cue EphrinB2 in the optic chiasm. Together, these findings position β2 laminins as a novel regulator of the ipsilateral projection.

## Introduction

Retinal ganglion cells (RGCs) collect visual information gathered by photoreceptors and transmit it to postsynaptic targets in the brain through their axons that collectively form the optic nerves and tracts. RGC axons enter the brain at the optic chiasm, a decision point in the mid-diencephalon where axons are routed to either the ipsilateral or contralateral side of the brain.

The optic chiasm is a critical point in the development of the optic pathway, where RGC axons selectively sort into the correct optic tract. In mice, axons from dorsonasal RGCs cross into the contralateral optic tract. In contrast, the axons from ventrotemporal RGCs do not cross and, instead, enter the ipsilateral tract (Colello and Guillery, 1990; Drager and Olsen, 1980; Godement et al., 1990). This selective decussation of RGC axons ensures the orderly establishment of circuits in the brain of binocular animals.

The selective decussation of RGC axons is orchestrated by radial glia and midline neurons in the optic chiasm (Marcus et al., 1995; Sretavan et al., 1994). These cells express axon guidance molecules that direct axons during decussation. Radial glia express Sema6D, NrCAM, and PlexinA1 to attract axons into the contralateral tract, as well as EphrinB2 to repel non-crossing axons into the ipsilateral tract (Kuwajima et al., 2012; Williams et al., 2003). Midline neurons express NrCAM and PlexinA1, driving crossing axons into the contralateral tract (Kuwajima et al., 2012).

RGC axon-selective decussation occurs in three phases that extend from E12 to P0. Each phase is defined by the RGCs that extend axons to the optic chiasm at the time. The early phase (E12-14) is when the first axons from RGCs in the central retina reach the chiasm and most cross into the contralateral tract (Colello and Guillery, 1990; Godement et al., 1987; Marcus et al., 1995). In the peak phase (E14-E17), most of the axons that reach the chiasm are from dorsonasal ganglion cells that also cross into the contralateral tract (Colello and Guillery, 1990; Godement et al., 1987; Marcus et al., 1995). The remaining axons are from ventrotemporal RGCs; these do not cross as they are repelled into the ipsilateral tract (Godement et al., 1990; Godement et al., 1994; Guillery et al., 1995). During the late phase (E17-P0), axons from the last RGCs born in the ventrotemporal retina cross into the contralateral tract (Drager, 1985). Consequently, the timing of birth and the retinal location where RGCs are born and determine whether their axons enter the contralateral or ipsilateral tract.

Based on their birthplace in the retina, RGCs acquire a lateral identity conferred by specific transcription factors. Contralateral RGCs are specified by Pou3f1 and Isl2 (Fries et al., 2023; Kuwajima et al., 2012; Pak et al., 2004). These contralateral-specific transcription factors are expressed in the dorsonasal retina throughout the phases of RGC selective axon routing. Pou3f1 is expressed in most post-mitotic differentiating RGCs and in developing amacrine cells (Fries et al., 2023). Isl2 is expressed in most RGCs but is absent in the ventrotemporal RGCs that express Zic2. The SoxC transcription factors (Sox4, 11, and 12) do not specify contralateral RGCs but do promote their differentiation (Kuwajima et al., 2012). In contrast, ipsilateral RGCs in the ventrotemporal retina are specified by the transcription factor Zic2, which is expressed during the peak phase of RGC selective decussation (Herrera et al., 2003).

Genetic deletion of Pou3f1 or Isl2 results in a larger ipsilateral projection due to an increase in the number of ipsilateral RGCs that are Zic2 positive (Fries et al., 2023; Pak et al., 2004). This suggests that Pou3f1 and Isl2 specify contralateral RGCs by repression of the ipsilateral program. Knockdown of the SoxC transcription factors results in a reduced number of contralateral RGCs without a change in the number of Zic2 positive ipsilateral RGCs. This suggests that these transcription factors control the differentiation of contralateral RGCs through an independent genetic pathway from Pou3f1 and Isl2 that does not repress Zic2 (Kuwajima et al., 2012). In contrast to the genetic deletion of contralateral-specific transcription factors, decreased levels of Zic2 result in fewer ipsilateral RGCs, leading to a nearly absent ipsilateral projection (Herrera et al., 2003). In summary, Pou3f1, Isl2, the SoxC family, and Zic2 are the transcription factors that determine the laterality type of RGCs.

RGC selective axon decussation is also influenced by cues provided by the extracellular matrix (ECM). Hyaluronan, an abundant ECM protein, regulates RGC axon decussation, and disrupting its function decreases the size of the ipsilateral projection (Lin et al., 2007). Heparan sulfate proteoglycans (HSPGs) are also linked to RGC axon decussation. Genetic deletions of enzymes that generate the sulfate modifications on HSPGs result in axon defasciculation and abnormal axon routing (Conway et al., 2011; Pratt et al., 2006).

Other ECM proteins implicated in axon guidance are laminins. A zebrafish genetic screen to isolate axon guidance mutants found that mutations in Lama1, Lamb2, and Lamc1 genes resulted in the routing of RGC axons that normally project to the contralateral tectum in the ipsilateral side of the brain (Barresi et al., 2010; Karlstrom et al., 1996; Paulus and Halloran, 2006). Supporting the role of laminin in axon guidance, a canonical laminin receptor, Dystroglycan (DG), has also been implicated in RGC axon selective decussation (Clements and Wright, 2018). These findings demonstrate that the ECM and its receptors regulate the path of RGC axons during selective decussation.

Basement membranes are specialized sheets of ECM composed of a basic toolkit of laminins, collagen type IV, and heparan sulfate proteoglycans (HSPGs) (Hynes, 2012). Of these, laminins are indispensable because they nucleate basement membrane formation (McKee et al., 2007). Basement membranes regulate cell polarity, proliferation, differentiation, apoptosis, migration, adhesion, and cytoskeletal organization (Yurchenco, 2011). Laminins are integral proteins of basement membranes that are heterotrimeric molecules composed of an α, β, and γ chain. Five genes encode α chains (Lama1-5), three encode β chains (Lamb1-3), and three encode γ chains (Aumailley et al., 2005). Laminins containing the β2 subunit are abundant in the inner limiting membrane (ILM) and pial basement membrane (PBM) (Hunter et al., 1992; Libby et al., 2000; Radner et al., 2013).

We hypothesize that β2 laminins in the PBM of the optic chiasm are required for the selective decussation of RGC axons. Consistent with our hypothesis, we found that genetic deletion of β2 laminins increased the proportion of non-decussating axons routed into the ipsilateral tract. Loss of β2 laminins caused fragmentation of the PBM and changes to radial glia basal processes and endfeet elaboration. This loss resulted in reduced expression of the laminin receptors β1-Integrin and β-dystroglycan in radial glia. These changes in the specialized niche of the optic chiasm may affect the expression of axon guidance cues by radial glia.

We found that the expression of the ipsilateral axon guidance cue EphrinB2 persists beyond its normal developmental window, likely extending the time window for ipsilateral guidance into the late phase of RGC axon-selective decussation. We also found that the number of ipsilateral RGCs is increased during this late phase of axonal decussation. Together, these results suggest that β2 laminins control the size of the ipsilateral projection by regulating the generation of the correct number of ipsilateral RGCs and the time window of expression of EphrinB2 in the optic chiasm.

## Materials and methods

### Animals

C57BL/6J wild-type, Lamb2^-/-^ (Lamb2^tm1Jrs^), and Lamc3^-/-^ (Lamc3^tm1.1Wjbr^) mice were used in the studies. Deletion of the Lamb2 and Lamc3 genes and the generation of the knockout lines were previously described (Li et al., 2012; Noakes et al., 1995). The Lamb2 line was maintained as heterozygotes, and the Lamc3 line as homozygotes. The sex of the embryos was not assessed for the experiments. The morning that the vaginal plug was observed was considered E0.5. For embryo collection, pregnant dams were euthanized by CO2 asphyxiation. Embryos at a specified age were collected and genotyped, with wild-type (WT) littermates serving as controls. Postnatal animals were euthanized by transcardial perfusion. All animal procedures were approved by the Institutional Animal Care and Use Committee (IACUC) of Upstate Medical University and adhered to the National Institutes of Health *Guide for the Care and Use of Laboratory Animals*.

### Anterograde labeling of retinal projections and measurement

Heads from P4 animals were fixed in 4% paraformaldehyde (PFA) overnight at 4°C. The fixative was then removed by washing the heads in 1X phosphate-buffered saline (PBS, 137 mM sodium chloride, 10 mM sodium phosphate, pH 7.4) 3 times for 5 min each. A small incision was made at the ora serrata, and the cornea and iris were cut, circling the ora serrata, leaving a piece intact to create a cornea/iris flap. The flap was lifted, the lens removed, and a small crystal of DiI (DiIC18(3), Invitrogen) was placed at the optic nerve head. The lens was placed back in the eye, and the cornea/iris flap was placed back over the lens. The DiI-labeled heads were then incubated at 37°C for 14 days in 1X PBS containing 0.02% sodium azide. After 14 days, the heads were washed in 1X PBS 3 X 10 min, and the brains were removed from the skull with care not to damage the optic chiasm or tracts. The optic chiasm and projections in the intact brain were imaged and photographed in a Leica MZ16 fluorescent dissection microscope.

The proportion of DiI-labeled ipsilateral and contralateral projections was determined by image analysis in FIJI (Schindelin et al., 2012). The area of each projection was traced from the midline to the edge of the ventral thalamus and measured. The area of each projection was used to calculate its proportional area. The proportion was determined by dividing each region by the total measured area.

### Immunohistochemistry

The antibodies used are detailed in the Key Resources Table. Embryo heads were fixed in 4% PFA overnight at 4°C. Heads were then washed in 1X PBS 3 times for 5 minutes each, and cryoprotected using sucrose solutions of increasing concentration. Heads were placed in 5%, 10%, 20%, and 30% sucrose in 1X PBS overnight at 4°C or until the head sank in the solution. Half of the 30% sucrose solution was replaced with OCT medium, mixed thoroughly by stirring and inverting, and the heads were incubated in this mixture overnight at 4°C on a shaker. The heads were then embedded in OCT and kept at -80°C until sectioning. Cryostat sections of the optic chiasm or retina (20 µm) were obtained. The sections were washed in 1X PBS, pH 7.5, 3 times for 10 min each, and then incubated in blocking solution (5% Normal Donkey Serum (NDS) and 0.2% Triton X-100 in 1X PBS) for 1 hour at room temperature. The sections were then incubated in primary antibodies overnight at 4°C in antibody diluting solution (5% NDS, 0.2 % Triton X-100, 1:15,000 Hoechst), washed in 1X PBS 3 X 10 min, and incubated with secondary antibody in antibody diluting solution for 2 hours at room temperature. The slides were then washed in 1X PBS 3 times for 10 min each and coverslipped using Prolong Gold or Glass antifade mounting medium (Thermo Fisher). Antigen retrieval with sodium Citrate buffer (pH 6.0) was performed before the blocking step for immunolabeling with Zic2 and Brn3a antibodies (Slavi et al., 2023).

To detect basement membrane proteins and visualize radial glial endfeet, embryo heads were fixed in 0.05% PFA for 1 hour on ice. The sections obtained from these lightly fixed heads were kept at -20°C during collection and until staining. The sections were fixed in ice-cold methanol for 2 minutes, washed in 1X PBS pH 7.5 3 times for 10 min each, and then incubated in blocking solution (5% Normal Donkey Serum (NDS) and 0.05% Triton X-100 in 1X PBS) for 1 hour at room temperature. The sections were then incubated in primary antibodies overnight at 4°C in antibody diluting solution (5% NDS, 0.01% Triton X-100, 1:30,000 Hoechst), washed in 1X PBS 3 X 10 min, and incubated with secondary antibody in antibody diluting solution for 2 hours at room temperature. The slides were then washed in 1X PBS 3 times for 10 min each and coverslipped using Prolong Gold or Glass antifade mounting medium (Thermo Fisher).

### In situ hybridization

The EphrinB2 coding sequence was PCR-amplified from the cDNA clone BC057009 (Horizon Discovery; Lafayette, Colorado) after sequence verification. The first 905 bp of the coding sequence was amplified using primers 5’-ATGGCCATGGCCCGGTC-3’ and 5’-CGCTGTCTGCAGTCCTTAGTGG-3’ to obtain a similar probe as described previously (Birgbauer et al., 2000) but excluded the last 107 bp of the coding sequence that has an 84% identity (Clustal Omega alignment) to the 3’end of EphrinB1 and EphrinB3. PCR amplicons were TA-cloned into pGEMTEZ (Promega; Fitchburg, WI) and sequence-verified. Digoxigenin-labeled riboprobes were transcribed in vitro from pGEMTEZ.EphrinB2 (SpeI, T7 polymerase for antisense and PspMOI, SP6 polymerase for sense), using the T7 or SP6 HiScribe High Yield RNA Synthesis kits (New England Biolabs; Ipswich, MA). For *in situ* hybridization on tissue sections, embryonic heads were fixed in 4% PFA overnight, washed 3 times in 1X PBS for 5 minutes each, cryoprotected in 10%, 20%, and 30% sucrose, and embedded in OCT. Optic chiasm sections (20 µm) were obtained and stored at -20°C until use. To minimize signal variability of the colorimetric reaction, WT and Lamb2^-/-^ sections were placed next to each other on the same slide. In situ hybridization was performed as previously described (Martinez-De Luna et al., 2013). All sequences and plasmids are available upon request.

### Confocal microscopy and fluorescence intensity measurements

Confocal images were acquired in the Neuroscience Program Microscopy Core at Upstate Medical University. Optic chiasm and retina sections were imaged on a Zeiss LSM780 laser scanning confocal microscope, equipped with an Argon laser and a He/Ne 633nm laser, using a Zeiss Plan Apochromat 20X/0.8 NA dry and 63X/1.4 NA oil lens. Pinhole was set to 1AU with a resolution of 512 X 512 pixels. A line average of 16 was applied. Confocal Z-series were acquired using identical gain and laser power to image WT and Lamb2^-/-^ sections. Fluorescent intensity measurements of β1-Integrin, β-Dag, and EphrinB2 were performed using Fiji software. Measurements were acquired from WT and Lamb2^-/-^ optic chiasm (3-9 sections) with equal image threshold and background subtraction. The chiasm area was defined using Nestin staining to identify the anterior, middle, and posterior sections. The fluorescence of the blood vessels and axons were subtracted from the β1-Integrin and the β-Dag intensity measurements. Images of ipsilateral and contralateral RGC counts were acquired at 20X with a digital zoom of 1.5 and a 0.92 step size to differentiate cells from one another carefully. High-magnification, high-resolution images of the PBM and radial glia endfeet were obtained on a Leica SP8 at 100X and 1012 X 1012 pixels with a digital zoom of 1.5 using a HC PL APO 100x/1.4 Oil STED white lens.

### Blinding of the sample genotype for analysis

Fluorescence intensity measurements and cell counts were performed in a blinded manner with respect to the animals’ genotypes. In most cases, mages were acquired by someone other than the person analyzing them. After image acquisition, the Z-stacks were randomized using a Python script that assigned random labels to each file and generated a de-blinded list of the samples. The blinded Z-stacks were given to the person analyzing the images. The original identities of the images were revealed after the counts were performed. The Z-stack scrambler code can be found at https://github.com/Martinez-De-Luna-Lab/Sample-Randomizer.

### Statistical Analysis

The sample sizes are stated in the figures and captions. Comparisons between groups were performed using unpaired Student’s t-test and one-way analysis of variance

(ANOVA) with Dunnett’s multiple comparisons post-test as appropriate. Significance was set at p<0.05. Graph Pad Prism software (GraphPad Software, La Jolla, CA) was used for all analyses.

## Results

### The β2 and γ3 laminin subunits are expressed in the optic chiasm at the time of selective axon decussation

The optic chiasm in the ventral brain is underlined by the pial basement membrane (PBM). β2 and γ3-containing laminins are abundant in the PBM (Hunter et al., 1992; Libby et al., 2000; Radner et al., 2013). Therefore, we decided to examine their distribution in the optic chiasm. We immunolabeled sections from E15.5 and E17.5 embryos using previously characterized β2- and γ3-chain-specific antibodies (Li et al., 2012; Sasaki et al., 2002). We selected E15.5 and E17.5 because these ages correspond to the peak and late phases of axon growth through the optic chiasm, respectively (Petros et al., 2008).

At E15.5 and E17.5, β2 laminin subunit immunoreactivity is robust in the PBM underlying the axons of the optic chiasm (Fig. 1A, B, D, and E), where they are co-expressed with Perlecan, another core basement membrane protein (Fig. 1C). We also find strong β2 laminin immunoreactivity in the vascular basement membrane of blood vessels that display CD31 immunoreactivity (Fig. 1A, B, D, E, and F; arrowheads). In addition, localization of β2 laminin is also observed in the apical surface of the third ventricle, the pia mater, the dura that surrounds the RGC axons, and cells within Rathke’s pouch, the presumptive anterior pituitary (Fig. 1A, B, D, and E, white arrow). The immunoreactivity observed with the β2-chain specific antibody is absent in the Lamb2^-/-^ optic chiasm (Supp. Fig. 1).

**Figure 1.**
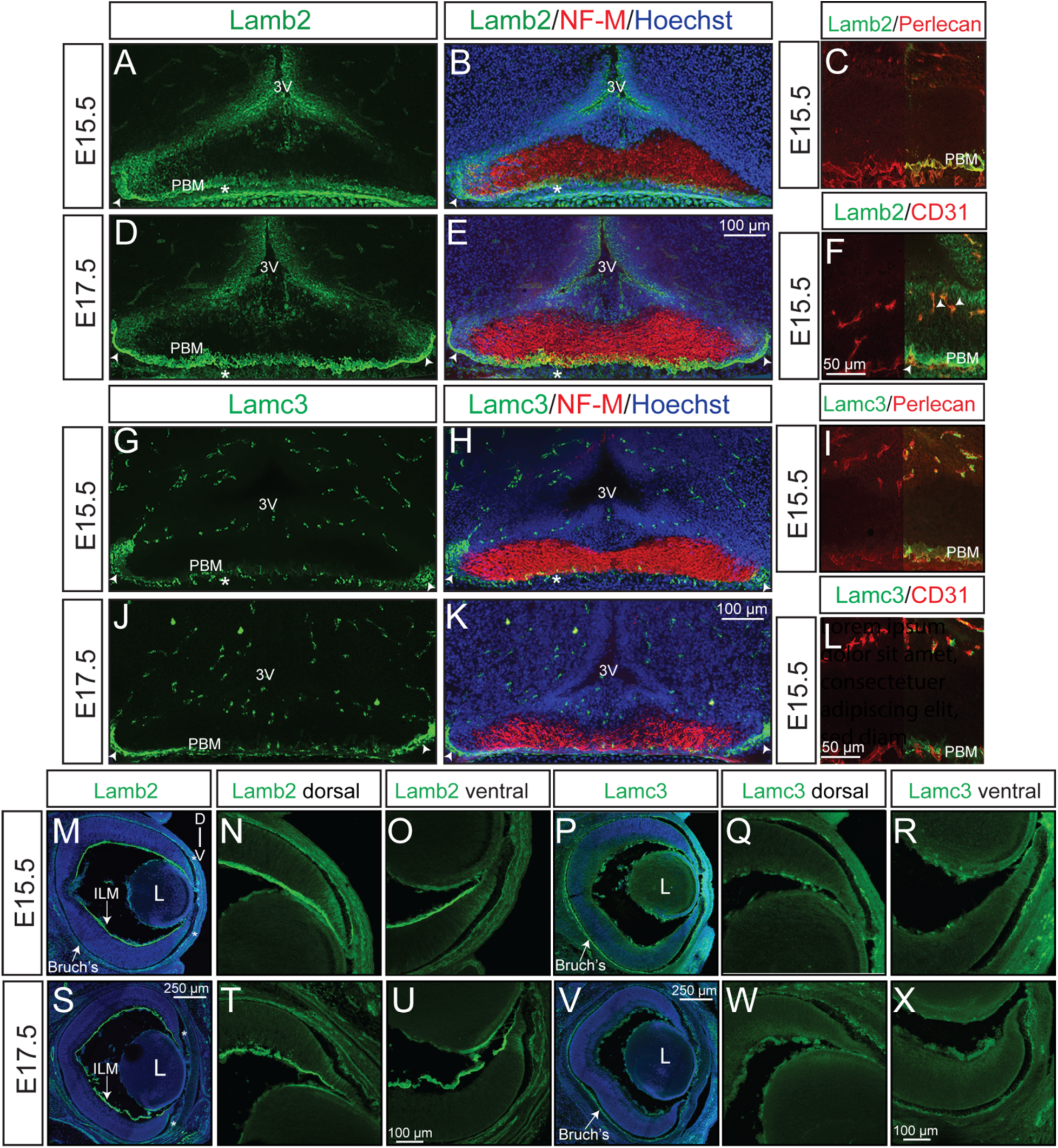
β2- and γ3-containing laminins are expressed in the optic chiasm and retina at the time of selective axon decussation. **(A, B, D, and E)** Lamb2 protein expression in the optic chiasm. Asterisk: pia mater; arrowhead: dura mater that covers the optic nerves and tracts. **(G, H, J, and K)** Lamc3 protein expression in the optic chiasm. Asterisk: pia mater; arrowhead: dura mater that covers the optic nerves and tracts. **(C, F, I, and L)** Confirmation of Lamb2 and Lamc3 expression in the PBM and microvasculature of the optic chiasm by co-staining with the PBM marker Perlecan and the endothelial cell marker CD31. **(M-X)** Lamb2 and Lamc3 protein expression in the retina with magnified views of the dorsal and ventral retina. D: dorsal; V: ventral. Asterisk: Retina periphery; L: lens; ILM: inner limiting membrane; Bruch’s: Bruch’s membrane. N=3 embryos analyzed for each panel.

We next investigated the protein distribution of the laminin γ3 subunit using a previously validated antibody (Li et al., 2012). The protein distribution of the laminin γ3 subunit is more restricted than that of the β2 subunit in the developing hypothalamus and optic chiasm. At E15.5 and E17.5, γ3 laminin is detected in the PBM where Perlecan is also present (Fig. 1G, H, J, K and I), but in lower abundance than β2 laminin. The laminin γ3 chain is a component of the CNS microvasculature (Li et al., 2012). Consistent with previous findings, we detected robust γ3 immunoreactivity in the basement membrane of the hypothalamic microvasculature and the pial vasculature, positive for CD31 (Fig. 1G, H, J, K and L). We also found strong γ3 laminin immunoreactivity in the dura that envelops the RGC axons and at the edges of the chiasm (Fig. 1G, H, J, and K; arrowheads).

We also determined the protein distribution for the β2 and γ3 laminin subunits in the retina. At E15.5, β2 laminins are abundant in the inner limiting membrane (ILM), the basement membrane of the retina (Fig. 1M-O). The ILM expression of β2 laminins extends into the dorsal and ventral ciliary marginal zone (CMZ) with some expression in the peripheral dorsal and ventral retina (Fig. 1N and O). In addition to the ILM and CMZ, β2 laminins are observed in Bruch’s membrane abutting the retinal pigment epithelium, the lens epithelium, and fragments of the hyaloid vasculature attached to the lens (Fig. 1M-O). At E17.5, the distribution of β2 laminins is very similar to E15.5 (Fig. 1S-U). However, at E17.5, although β2 laminins are expressed in the dorsal and ventral peripheral retina, the expression becomes more enriched in the ventral peripheral retina and CMZ (Fig. 1S and U).

In contrast to β2 laminin expression, at E15.5 and E17.5, the expression of γ3 laminins is low in the ILM and high in Bruch’s membrane, a finding that is consistent with previous work (Fig. 1P and V) (Li et al., 2012). We did observe very faint expression of γ3 laminins in the dorsal and ventral ciliary marginal zone at E15.5 (Fig. 1Q and R). The expression of γ3 laminins was absent from the ciliary marginal zones at E17.5 (Fig 1W and X). We also observed γ3 laminin expression in the fragments of the hyaloid vasculature attached to the ILM and lens at both ages (Fig. 1P-R and V-X).

Together, these results demonstrate that the β2 and γ3 laminin subunits are expressed in the developing optic chiasm. In the retina, β2 is predominantly expressed in the ILM, with early expression in the peripheral retina that then accumulates in the ventral CMZ. γ3 laminins, in contrast, are predominantly expressed in Bruch’s membrane at the ages examined.

### The selective decussation of ipsilateral RGC axons is dependent on laminins

Laminins have been implicated in the guidance of axons in commissures of the central nervous system and the optic chiasm in zebrafish (Barresi et al., 2010; Paulus and Halloran, 2006). Based on these studies and the expression patterns we observed in the optic chiasm, we hypothesized that β2 and γ3 laminins in the PBM are positioned to provide cues that influence the selective decussation of retinal axons. To test this hypothesis, we asked if the selective decussation of RGC axons was altered in Lamb2 and Lamc3 knockout mice compared to WT controls. We anterogradely labeled RGC axons from the right eye with DiI to measure the area of the retinal projections (Figure 2A). We used these measurements to calculate the proportional area of the ipsilateral and contralateral optic tracts in WT, Lamb2^-/-^, and Lamc3^-/-^ mice (Fig. 2A).

**Figure 2.**
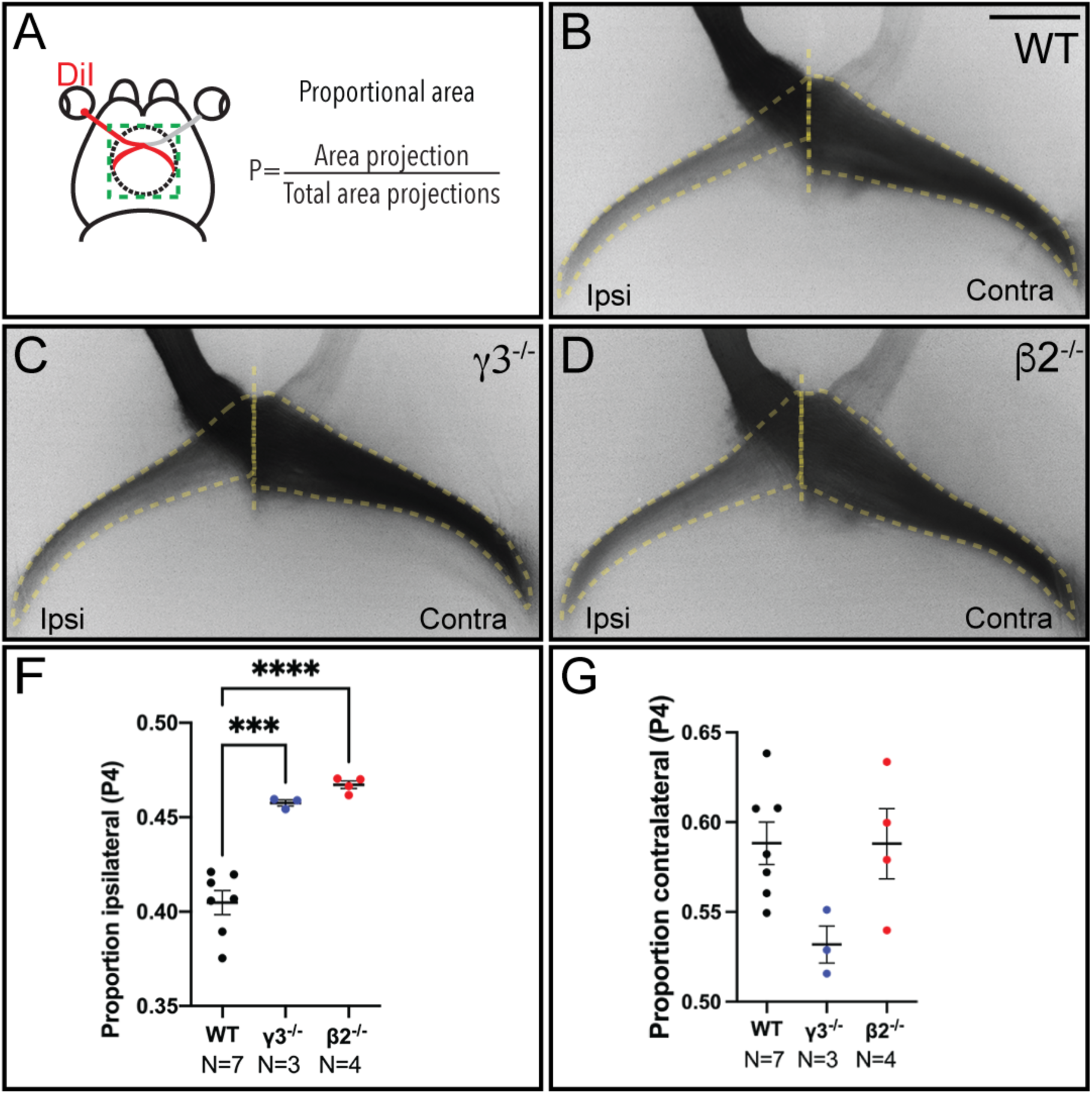
The proportion of the ipsilateral projection is increased in Lamc3^-/-^ and Lamb2^-/-^ mice. **(A)** Experimental schematic showing placement of DiI crystals and anterograde labeling of retinal projections. The proportional areas of the projections were calculated using the formula. **(B-D)** Labeled RGC projections in the WT, Lamc3^-/-^, and Lamb2^-/-^ mice. Dashed lines trace the area of each projection. The perpendicular line marks the optic chiasm midline. **(F)** The proportional area of the ipsilateral projection. **(G)** The proportional area of the contralateral projection. Statistical significance was calculated using a One-way ANOVA with Dunnett’s multiple comparisons post-test. Error bars represent the S.E.M. N=number of mice.

In P4 WT mice, the proportional area of the ipsilateral projection was 0.40 ± 0.016 of the total size of the projections, while the proportional area of the contralateral projection was 0.60 ± 0.012 (Fig. 2B, F, and G). We found that in the Lamc3^-/-^ and Lamb2^-/-^ mice, the proportion of the ipsilateral projection was increased compared to wild-type (Lamc3^-/-^: 0.46 ± 0.001, *p*=0.0002; Lamb2^-/-:^ 0.47 ± .002, *p*<0.0001) (Fig. 2C, D, and F). The size of the contralateral projection was not changed in the Lamc3^-/-^ and Lamb2^-/-^ mice (Lamc3^-/-^: 0.53 ± 0.01, p=0.056; Lamb2^-/-^: 0.58 ± 0.02, p= 0.059) (Fig. 2C, D and G).

These results suggest that β2 and γ3-containing laminins present in the PBM contribute to the regulation of the selective decussation of RGC axons.

### β2 laminins are necessary for the integrity of the PBM and the normal morphology of radial glia basal processes and endfeet

The midline radial glia that control RGC axonal decussation have endfeet that contact the PBM (Colello and Guillery, 1992; Marcus et al., 1995). In the developing cortex, loss of β2 laminins resulted in dysmorphic radial glia with misaligned, highly branched basal processes and poorly formed endfeet (Radner et al., 2013). Together with these defects in the radial glia, the absence of β2 laminins resulted in fragmentation of the inner limiting membrane (ILM) in the retina and the PBM in the cortex (Pinzon-Duarte et al., 2010; Radner et al., 2013).

Based on these findings, we determined if similar defects were present in the PBM and radial glia of the optic chiasm. We asked if loss of β2 laminins compromised the integrity of the PBM and disrupted the morphology of the midline radial glia. We immunolabeled optic chiasm sections from WT and Lamb2^-/-^ embryos at E15.5 and E17.5 to examine the integrity of the PBM and the shape of the radial glia basal processes and endfeet.

At E15.5, the WT PBM is a continuous sheet of Nidogen immunoreactivity at the pial surface (Fig. 3A). The radial glia basal processes extend, parallel to each other, to contact the PBM with their endfeet (Fig. 3B and C). In contrast, in the Lamb2^-/-^ optic chiasm, the PBM is fractured and only apparent in patches (Fig. 3G). Below the fragmented PBM, the pial blood vessels are prominent, as shown by Nidogen immunostaining (Fig. 3G). The radial glial basal processes are disorganized and do not run parallel to one another toward the PBM. Instead, the basal processes are undulated and overlap with neighboring basal processes (Fig. 3H). The basal processes also appear to lack defined endfeet and attach to fragments of the PBM and the pial blood vessels (Fig. 3H and I).

**Figure 3.**
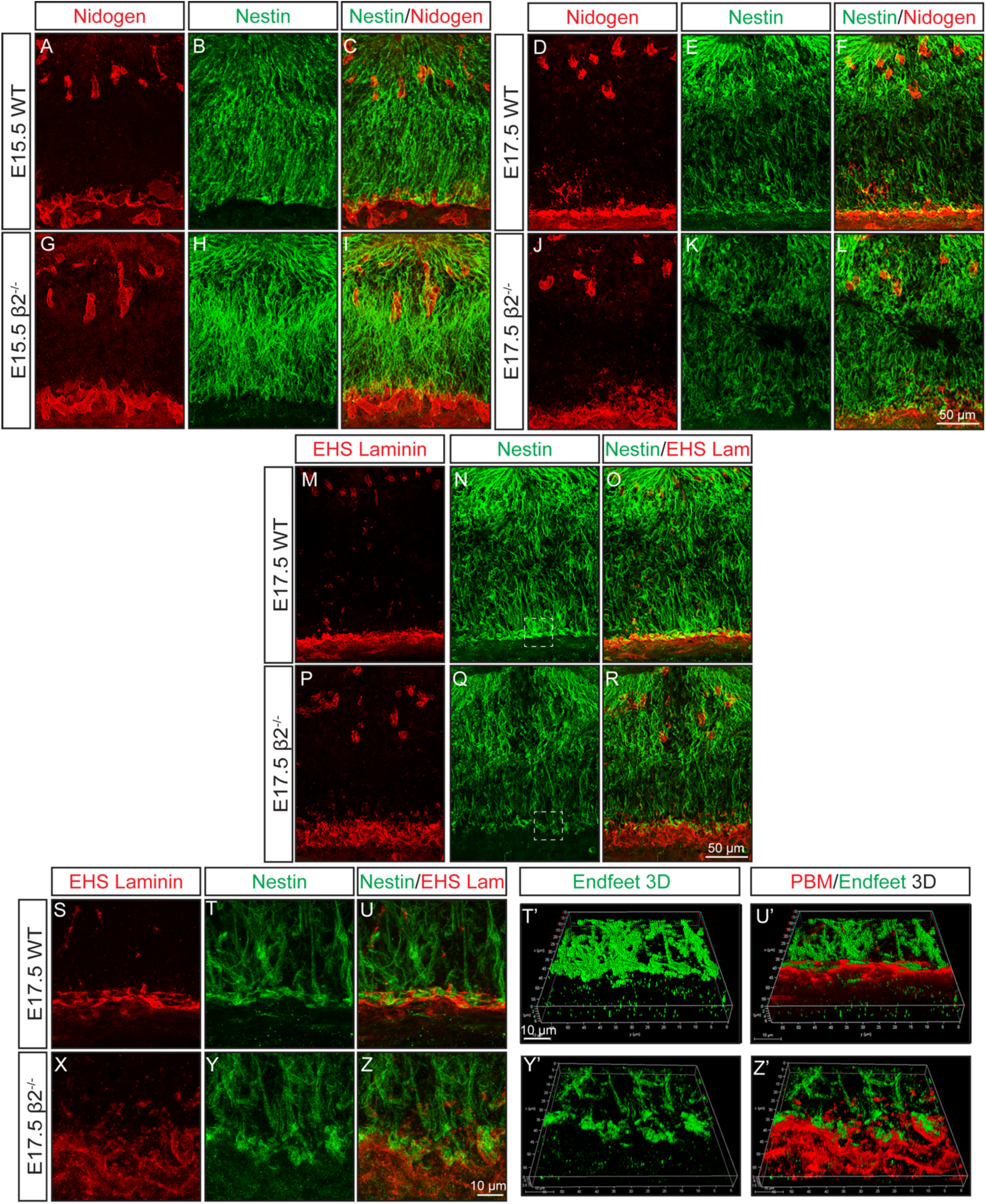
Loss of β2-containing laminins leads to fragmentation of the pial basement membrane and radial glia with abnormal basal processes and endfeet. (**A-R**) Organization of the PBM (red) and radial glia basal processes (green) at E15.5 and E17.5. The PBM fragmentation at E17.5 is shown using Nidogen and EHS Laminin. **(S-Z)** Magnified and higher resolution view of the dashed square in panels N and U showing the fragmentation of the PBM and disrupted elaboration of the radial glia endfeet in the Lamb2^-/-^ chiasm. **(T’, U’, Y’, Z’)** 3D-volume reconstruction from confocal Z-stacks in T, U, Y and Z. N=3 embryos per experiment at each age.

At E17.5, the WT PBM is a continuous, thicker sheet of Nidogen and EHS Laminin immunoreactivity at the pial surface (Fig. 3D and M). The WT radial glia basal processes are elongated and organized nearly parallel to one another (Fig. 3E and N). The basal processes terminate in mature, mallet-shaped endfeet that attach to the PBM (Fig. 3E, F, N, and Q). In contrast, in the Lamb2^-/-^ optic chiasm, the PBM is fragmented and lacks the continuity of Nidogen and EHS Laminin immunoreactivity in the WT (Fig. 3J and P). Also, blood vessels in the pia mater under the PBM exhibit increased Nidogen and EHS Laminin immunoreactivity, giving the pia mater a ruffled appearance (Fig. 3J and P). The radial glial basal processes are thinner and less parallel to one another, with underdeveloped endfeet that lack their typical mallet-shaped morphology (Fig. 3K and Q). The abnormally shaped endfeet extend past the fragmented PBM to bind to pial cells and possibly blood vessels (Fig. 3L and R).

Next, we further investigated the fragmentation of the PBM and the elaboration of endfeet. We acquired higher-magnification, higher-resolution images of the radial glial endfeet and PBM in the same sections stained with EHS Laminin at E17.5 (Fig. 3S-Z). We then used these images to generate 3D-reconstructions to better visualize the endfeet and PBM (Fig. 3T’, U’, Y’ and Z’). At higher magnification and resolution, the PBM is a continuous sheet in which the mallet-shaped endfeet are embedded (3M-O). The radial glia endfeet attach to the PBM in close apposition to one another, forming the glia limitans (Fig. 3T, U, T’ and U’). The PBM in the Lamb2^-/-^ chiasm is fragmented and nearly absent (3X, Z and Z’). The radial glia endfeet in the Lamb2^-/-^ chiasm terminate in abnormally shaped endfeet that fasciculate with each other to form clumps (Fig. 3Y, Z, Y’ and Z’). These clumped endfeet do not form a continuous glia limitans as in the WT (Fig.3Y and Y’). Despite their abnormal morphology, the endfeet of the Lamb2^-/-^ radial glia attach to remnants of the PBM and occasionally to blood vessels in the pia mater (Fig. 3X, Z and Z’).

In summary, in the absence of β2 laminins, the PBM in the optic chiasm becomes fragmented, and the radial glia become dysmorphic and have poorly elaborated endfeet. Together, these findings suggest that the organization of the PBM and the radial glia that direct axon guidance is dependent on β2 laminins.

### The expression of laminin receptors in the radial glia is reduced in the Lamb2^-/-^ optic chiasm

The disruption of the PBM and changes to the radial glia basal processes and endfeet led us to investigate if the expression of the laminin receptors β1-Integrin and β-Dystroglycan (β-Dag) was reduced in the Lamb2^-/-^ optic chiasm compared to the WT.

Radial glia express β1-integrin and Dag (Loulier et al., 2009; Myshrall et al., 2012). Conditional deletion of these receptors in radial glia results in fragmentation of the PBM and dysmorphic radial glial basal processes that detach from the PBM (Graus-Porta et al., 2001; Myshrall et al., 2012; Satz et al., 2010). Given the similar changes in the PBM and radial glia in Lamb2^-/-^ embryos and because laminins regulate the expression of their own receptors (Condic and Letourneau, 1997), we asked if deletion of Lamb2 affected β1-Integrin and β-Dag expression in the radial glia of the optic chiasm.

The radial glia in the WT optic chiasm expressed β-Dag and β1-Integrin with strong immunolabel detected in the ventricular zone surrounding the radial glia cell bodies and in the glial endfeet that abut the PBM (Fig. 4A-D). We also detected β-Dag and β1-integrin expression in the blood vessels and axons, which were excluded from our measurements (Fig. 4A-D). In the Lamb2^-/-^ optic chiasm, β-Dag and β1-integrin expression was reduced in the radial glia cell bodies, processes, and endfeet as well as the blood vessels and axons (Fig. 4F-I; N=4). We found that the reduction in both receptors is a modest 20% but significantly different from the WT (Fig. 4E and J; N=4; WT: 100% and Lamb2^-/-^: 79.74 ± 2.27% for β1-Integrin and WT: 100% and Lamb2^-/-^: 81.46 ± 2.09% for β-Dag). These results show that genetic deletion of β2 laminins reduces the expression of two canonical laminin receptors in the radial glia of the optic chiasm and suggest that the changes in Lamb2^-/-^ radial glia are due to the reduction in expression of these receptors.

**Figure 4.**
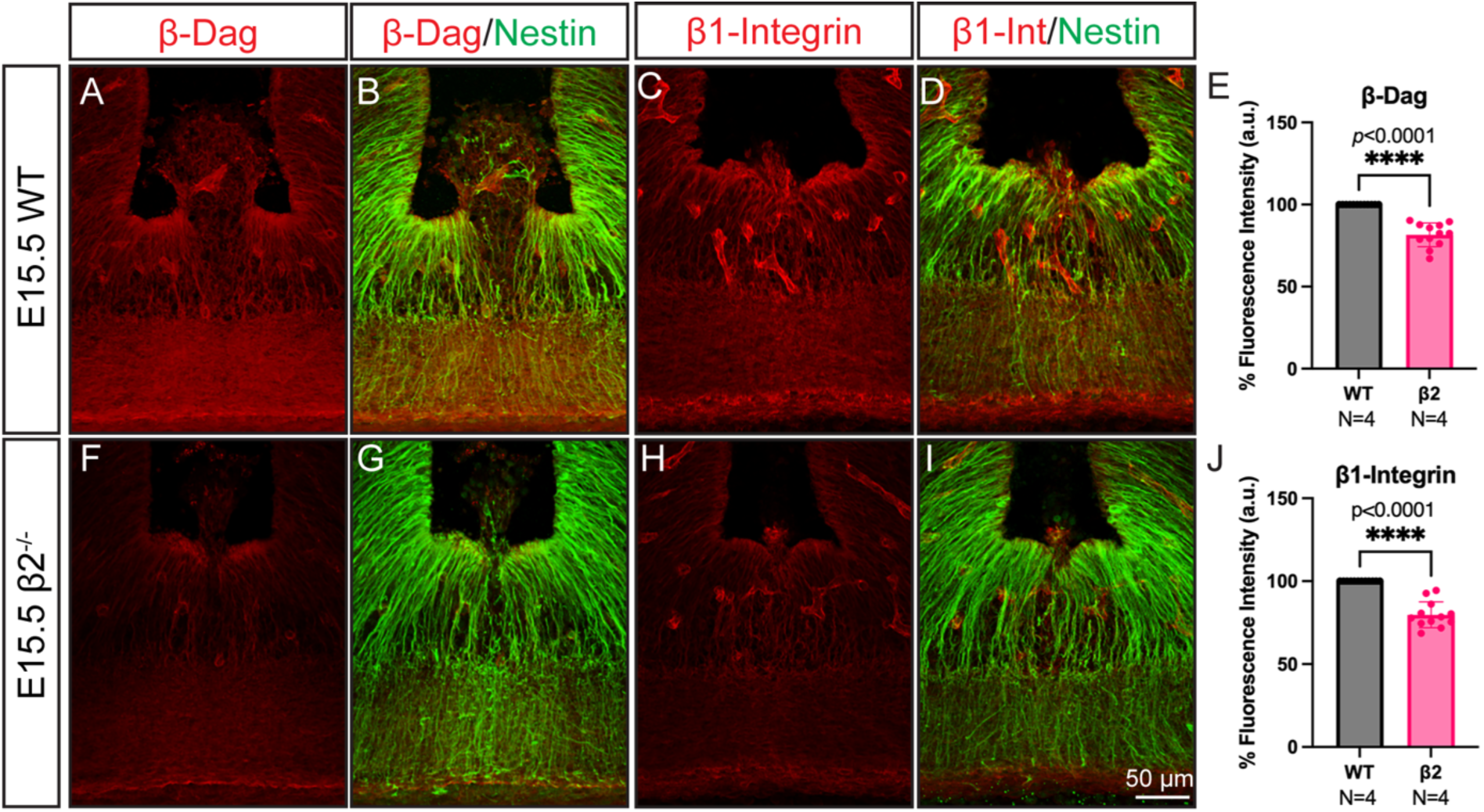
β1-Integrin and β-Dag expression is reduced in the absence of β2 laminins. **(A, B, and F, G)** Comparison of β-Dag protein expression (red) in WT and Lamb2-/- radial glia (green). **(C, D, and H, I)** Comparison of β1-Integrin expression (red) in WT and Lamb2^-/-^ radial glia (green). **(E)** Quantification of the percent of β-Dag fluorescence intensity in the radial glia. **(J)** Quantification of the percent of β1-Integrin fluorescence intensity in the radial glia. N=number of mice.

### EphrinB2 expression persists into the late phase of retinal ganglion cell axon decussation in the absence of β2 laminins

EphrinB2 is expressed in radial glia, and its peak of expression is at E15.5 when most of the ipsilateral and contralateral RGCs are extending axons to the optic chiasm (Williams et al., 2003). EphrinB2 expression is reduced by E17.5, the late phase of axon decussation, when the remaining RGC axons entering the optic chiasm project contralaterally (Williams et al., 2003). We found that the size of the ipsilateral projection is increased in the Lamb2^-/-^ mice at P4 and that this change is accompanied by changes in the radial glia of the optic chiasm at the time of RGC selective decussation. These findings prompted us to ask if the expression of the axon guidance cue EphrinB2 was altered in the Lamb2^-/-^ radial glia.

We found that EphrinB2 expression was barely detectable in the E17.5 RC2-positive radial glia of the WT chiasm, with low levels in the basal processes as expected (Fig. 5A-C and G; N=4). This result is consistent with previous findings that EphrinB2 expression diminishes in the chiasm during the late phase of RGC axonal decussation (Williams et al., 2003). Surprisingly, in the Lamb2^-/-^ chiasm, we found that high EphrinB2 expression persisted in the RC2 positive radial glia into the late phase of axon decussation and that it was 70% higher than in the WT (WT: 100% and Lamb2^-/-^: 170 ± 9.86%; Fig. 5D-G; N=4). These results suggest that β2 laminins provide cues to the radial glia that control the expression of EphrinB2 during the shift from the peak to the late phase of RGC axon guidance in the optic chiasm.

**Figure 5.**
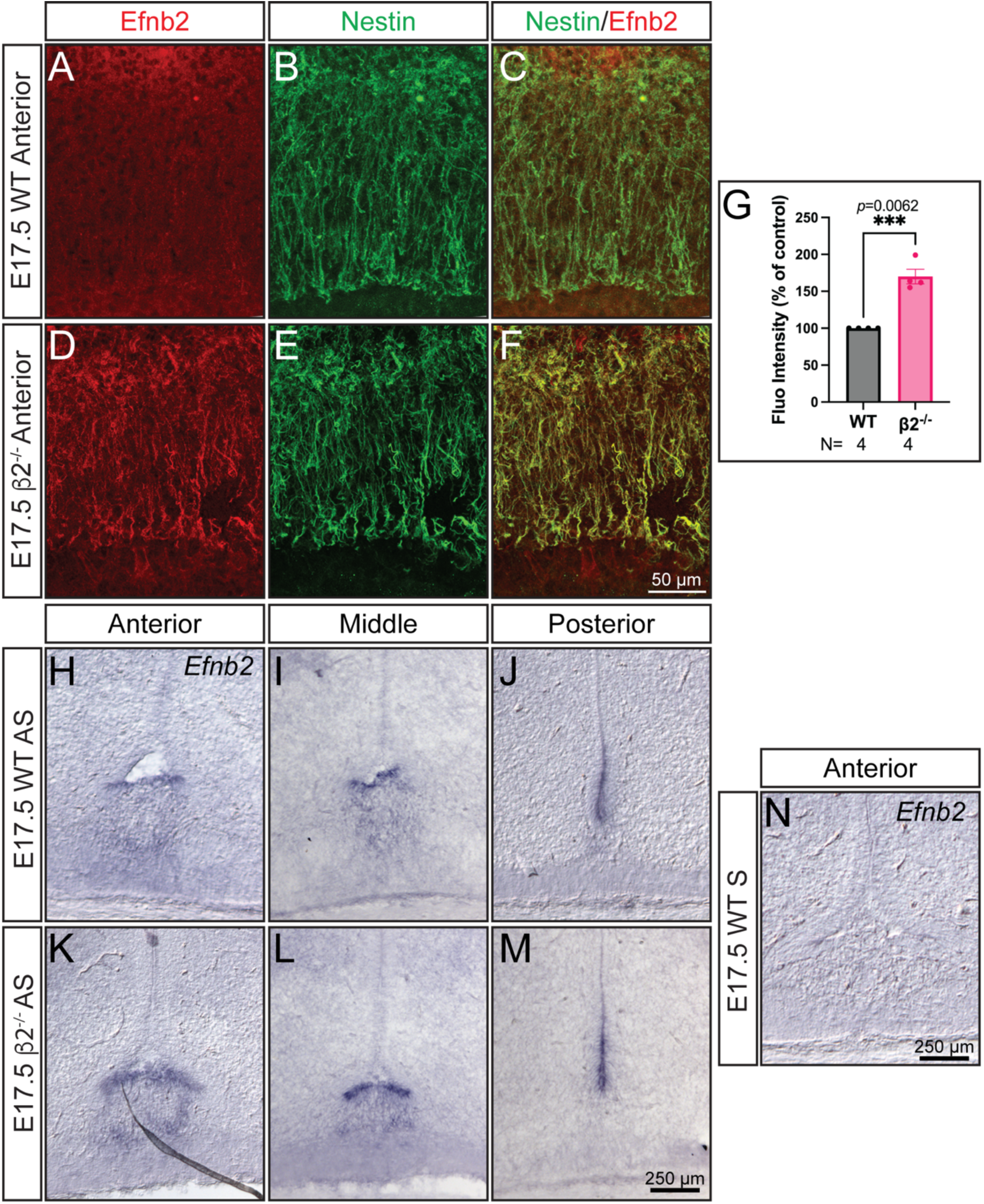
EphrinB2 expression persists in the radial glia during the late phase of RGC axon decussation. **(A-F)** EphrinB2 protein expression (red) in WT and Lamb2-/- radial glia (green) at E17.5, which corresponds to the late phase of RGC axonal decussation. N=number of mice. **(D)** Quantification of EphrinB2 fluorescence intensity in WT and Lamb2^-/-^ radial glia. **(H-M)** EphrinB2 mRNA expression in the optic chiasm with antisense RNA probe. **(N)** Lack of EphrinB2 expression with the control sense RNA probe.

Next, we performed in situ hybridization for EphrinB2 in optic chiasm sections from WT and Lamb2^-/-^ mice at E17.5 to confirm our immunolabeling results and determine whether EphrinB2 persistence was at the transcript level. In the anterior and mid WT chiasm, we detected the EphrinB2 transcript in the ventricular zone where the radial glia nuclei are located and in the processes of the radial glia extending into the RGC axonal tract (Fig. 5H and I). In the posterior WT chiasm, EphrinB2 transcripts were concentrated in the ventricular zone (Fig. 5J). In the Lamb2^-/-^ chiasm, EphrinB2 was also expressed in the cell bodies and processes of the radial glia in the anterior and mid chiasm, but the signal was more robust than in the WT (5K and L). Posteriorly, EphrinB2 expression was restricted to the ventricular zone and appeared more intense than in the WT (Fig. 5M). The sense probe derived from the same sequence as the antisense probe did not detect any EphrinB2 transcript in the sections (Fig. 5N). These results support the increased EphrinB2 protein distribution observed by immunolabeling and suggest that β2 laminins transcriptionally regulate the expression of EphrinB2 in the radial glia.

### The number of Zic2-positive retinal ganglion cells is increased in the absence of β2 laminins

Zic2 specifies the ventrotemporal RGCs that project ipsilaterally, and its expression is restricted to the peak phase of axon segregation (Herrera et al., 2003). We considered that the increased proportion of ipsilateral axons in the Lamb2^-/-^ mutants could result from an increase in the number of ipsilateral RGCs. We tested this possibility by immunolabeling and counting the number of Zic2-positive ipsilateral RGCs, Brn3a-positive contralateral RGCs, and Zic2 plus Brn3a double-positive cells at E15.5 and E17.5 in the ventrotemporal retina (Quina et al., 2005; Xiang et al., 1995). The counts were performed double-blind to the animals’ genotype.

At E15.5, the peak phase of RGC birth and axon segregation, the number of Zic2 (WT: 29.13±2.242 vs Lamb2^-/-^: 26.2±3.322; p=0.3670), Brn3a (WT: 13.62±1.578 vs Lamb2^-/-^: 10.87±0.8338; p=0.1355) and Zic2 plus Brn3a positive cells (WT: 3.43±0.6160 vs Lamb2^-/-^: 2.94±0.3025; p=0.4814) in the ventrotemporal retina of the Lamb2 knockout is the same as WT (Fig. 6A-I). We did not find a significant difference in the number of nuclei in the differentiated cell layer of the ventrotemporal retina (Fig. 6J) (WT: 95.68±2.75 vs Lamb2^-/-^: 99.42±1.528; p=0.2478).

**Figure 6.**
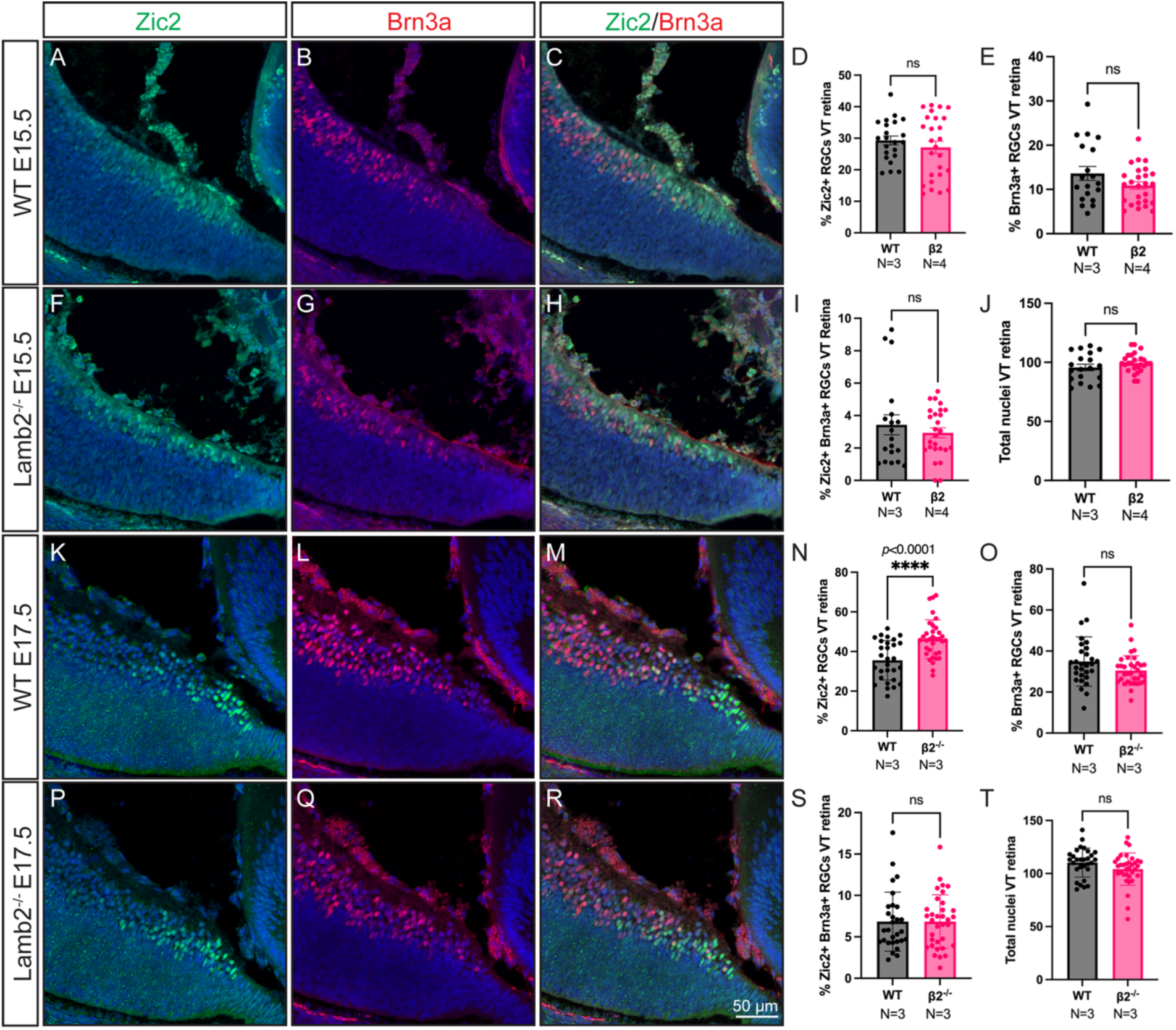
The number of ipsilateral RGCs is increased in the Lamb2^-/-^ ventrotemporal retina. **(A-C and F-H)** Zic2 positive ipsilateral (green) and Brn3a positive contralateral RGCs (red) in the ventrotemporal retina at E15.5. Nuclei are labeled with Hoechst (blue). **(K-M and P-R)** Zic2 (green) and Brn3a (red) RGCs in the ventrotemporal retina at E17.5. **(D and N)** Quantification of Zic2-positive cells. **(E and Q)** Quantification of Brn3a-positive cells. **(I and S)** Quantification of Zic2 and Brn3a double-positive cells. **(J and T)** Quantification of Hoechst-labeled nuclei in the differentiated cell layer of the ventrotemporal retina. N=number of mice.

Next, we examined the ventrotemporal retina at E17.5, when the last RGCs born in this region project contralaterally. Surprisingly, the number of Zic2 positive RGCs is significantly increased in the Lamb2^-/-^ ventrotemporal retina compared to the WT (WT: 35±1.861 vs Lamb2^-/-^: 46±1.68; p<0.0001) (Fig. 6K, N, P and S). In contrast, the percentage of Brn3a (WT: 34.87±2.230 vs Lamb2^-/-^: 30.49±1.241; p=0.093) or Brn3a/Zic2 (WT: 6.835±0.6604 vs Lamb2^-/-^: 6.849 ±0.5542; p=0.9875) double positive cells was not increased (Fig. 6L, M, O, Q, R and S). Additionally, as at E17.5, the total number of nuclei in the ventrotemporal retina did not change in the Lamb2^-/-^ retinas compared to the WT (WT: 110.3±2.547 vs Lamb2^-/-^: 104.2±2.615; p=0.0999) (Fig. 6T). These results suggest that β2 laminins control the number of ipsilateral RGCs in the ventrotemporal retina during the establishment of the ipsilateral projection.

## Discussion

We provide evidence that β2 laminins control the size of the ipsilateral projection during optic pathway formation. We demonstrate that β2 laminins in the optic chiasm maintain the integrity of the PBM and the organization of the radial glia scaffold that directs selective RGC decussation. We also show that β2 laminins provide cues to the radial glia that control the developmental expression of EphrinB2. Our findings suggest that β2 laminins have an active role in limiting EphrinB2 expression to its developmental expression window that begins in the peak phase and ends in the late phase of axonal decussation. We further show that β2 laminins function in the retina to control the number of ipsilateral RGCs in the developing ventrotemporal retina. Based on these findings, we propose that β2 laminins regulate RGC selective axon decussation by providing cues that control the number of ipsilateral RGCs and the timely expression of the ipsilateral cue EphrinB2 in the optic chiasm.

### β2 laminins in the organization of the optic chiasm

Radial glia in the developing brain are long, bipolar cells whose apical ends contact the ventricles and whose basal endfeet attach to the PBM. In the cortex, this bipolar shape is essential because newborn neurons use radial glia as guides during migration to the correct layer (Rakic, 1972). This guiding role depends on interactions with the ECM and related signaling. When PBM components such as β2 laminins, ECM receptors, or their downstream effectors are removed, the PBM breaks down, radial glia endfeet detach, and their basal processes become disrupted (Graus-Porta et al., 2001; Halfter et al., 2002; Myshrall et al., 2012; Radner et al., 2013; Satz et al., 2010). These defects show that ECM–receptor pathways are needed to maintain the PBM and the radial glia scaffold, and that radial glia function in cortical development relies on ECM interactions.

The radial glia of the optic chiasm share the stereotypical bipolar organization of their cortical counterparts, including their attachment to the PBM (Guillery et al., 1995; Marcus et al., 1995; Sretavan et al., 1994). However, the function of the radial glia in the optic chiasm is to direct growing RGC axons into the correct optic tract through the expression of axon guidance molecules on their cell surface (Kuwajima et al., 2012; Williams et al., 2006; Williams et al., 2003). RGC growth cones directly contact the radial glia basal processes in the chiasm while deciding whether to enter the ipsilateral or contralateral optic tract (Marcus et al., 1995). This makes the organization of radial glial basal processes critical for steering RGC growth cones through the glial scaffold in the optic chiasm.

Our finding that genetic deletion of β2 laminins increased the proportion of ipsilateral axons led us to investigate whether there were changes in the PBM and radial glia. Indeed, in the Lamb2^-/-^ optic chiasm, the PBM had compromised integrity, and the radial glia had abnormal basal processes with poorly elaborated endfeet. The basal processes frequently lacked attachment to the remaining fragments of PBM or terminated early before reaching the PBM or went past it. These changes strongly suggest that, like in the cortex, β2 laminins are necessary for maintaining the integrity of the PBM and for the organization of the radial glia basal processes in the optic chiasm.

The PBM’s fragmentation appeared progressive. In the peak phase of axonal decussation, the PBM was ruptured, but the PBM appeared more fragmented in the late phase. We observed progressive fragmentation with antibodies to EHS laminin and Nidogen, suggesting that the fragmentation was not due to changes in the expression of specific basement membrane proteins. The PBM fragmentation is likely due to the absence of secretion of laminin trimers containing β2 laminin. The analysis of β2 laminin expression showed that these laminins are expressed in the pia and ventricular zone of the ventricle. The pia mater contains the meninges, which are a significant source of basement membrane components that maintain the structure of the PBM (Beggs et al., 2003). It is likely that disruption of β2 laminin secretion from the pia in the β2 laminin knockout is one contributing factor to the loss of PBM integrity.

### Regulation of EphrinB2 expression by β2 laminins

Concurrent with the changes in the PBM and in the radial glia basal processes and endfeet, we found that EphrinB2 expression persisted beyond its normal developmental window in the Lamb2^-/-^ radial glia. The peak of EphrinB2 expression is during the peak phase of axon decussation when the ipsilateral projection is established. EphrinB2 expression is downregulated in radial glia at the initiation of the late phase of axon decussation, when the last RGCs born in the VT project contralaterally (Williams et al., 2003). Our findings demonstrate that β2 laminins function in the molecular pathway required to turn off EphrinB2 expression in the radial glia by the late phase of selective axon decussation.

The persistent expression of EphrinB2 in the late phase of axon decussation suggests that laminins regulate EphrinB2 expression through an adhesion-dependent mechanism that involves radial glia attachment to laminins via a canonical laminin receptor or other type of receptor. Laminins are known to regulate EphrinB2 expression in the developing retinal vasculature. Vessels in the retinal vasculature acquire either a venous or arterial fate. Laminin γ3 is expressed in the venous and arterial BM, but its receptor, Dag, is only expressed in arterial endothelial cells. Dag-laminin signaling promotes arterial fate by inducing EphrinB2 expression in arteries (Biswas et al., 2018). In contrast, the absence of Dag expression in venous endothelial cells leads to induction of the venous fate by activating Notch and upregulating EphB4 (Biswas et al., 2018). These previous findings support our hypothesis that laminins can regulate EphrinB2 expression.

In the optic chiasm, radial glia attach to the PBM and could receive signals from β2 laminins via either Dag or integrins, since both are expressed in these cells (Graus-Porta et al., 2001; Myshrall et al., 2012) and this study). The control of EphrinB2 expression could be mediated through Dag signaling, as in the retinal vasculature. However, laminin signaling through integrins is a plausible alternative. Ephrins have been implicated in mediating cell adhesion and cytoskeletal re-arrangements by increasing integrin signaling pathways (Davy and Robbins, 2000; Huynh-Do et al., 2002). In future studies, we will dissect the roles of the two major types of laminin receptors in regulating EphrinB2.

### β2 laminins in the generation of ipsilateral RGCs

Our study also identifies β2 laminins as a novel player in the acquisition of laterality in RGCs. Our data shows that β2 laminins restrict the number of ipsilateral RGCs in the VT retina during the late phase of RGC birth and axonal decussation. Genetic deletion of β2 laminins increased the number of Zic2-positive ipsilateral RGCs, suggesting that these laminins maintain the correct number of ipsilateral RGCs generated in the VT retina.

The specification of most ipsilateral RGCs by Zic2 takes place during the peak of RGC birth between E14 and E17 (Petros et al., 2008). During this developmental time window, Zic2 is expressed in the VT retina, where ipsilateral RGCs are found in the mouse (Herrera et al., 2003). After E17, Zic2 is downregulated in VT RGCs, and the last RGCs born in the VT retina acquire a contralateral fate (Drager, 1985; Drager and Olsen, 1980; Williams et al., 2006). β2 laminins are abundantly expressed in the ILM (Hunter et al., 1992; Libby et al., 2000). Our findings show that by E17.5, β2 laminins become enriched in the peripheral VT retina and ciliary marginal zone (CMZ). This finding suggests that β2 laminins are positioned to regulate the number of RGCs generated in the VT retina.

We observed an increased number of ipsilateral RGCs without a change in the total number of nuclei in the differentiated cell layer at E17.5. This finding suggests that β2 laminins may control the number of ipsilateral and contralateral RGCs in the VT retina by regulating cell fate. β2 laminins have an established role in regulating cell fate in the postnatal mouse retina (Serjanov et al., 2018). In the Lamb2^-/-^ retina, rods are overproduced at the expense of later-born bipolar cells and Müller glia (Serjanov et al., 2018). The change in cell fate was driven by a loss of retinal progenitor cell basal processes, which then altered the plane of cell division and changed the fate of the retinal cells generated. Specifically, rods were overproduced at the expense of the later-born bipolar cells and Müller glia (Serjanov et al., 2018).

The involvement of β2 laminins in the regulation of RGC laterality suggests that they may provide cues that influence the genetic cascade that switches the fate of RGCs in the VT retina from ipsilateral to contralateral during the late phase of RGC birth. Pou3f1 and Isl2 control the contralateral RGC fate. Genetic deletion of Pou3f1 or Isl2 results in an increased number of Zic2-positive ipsilateral RGCs (Fries et al., 2023; Pak et al., 2004). Furthermore, Pou3f1 directly binds the Zic2 promoter and represses its expression (Fries et al., 2023). β2 laminins may promote the expression of Pou3f1 or Isl2 in late-born RGCs. Future studies will determine if Zic2 RGCs are produced at the expense of Pou3f1 or Isl2-expressing contralateral RGCs. This would determine if β2 laminins provide cues to RPCs in the VT retina that control the lateral fate of RGCs.

The receptor and intracellular pathways through which laminin β2 laminins exert their function on RGC lateral fate are unknown. Interestingly, ipsilateral RGCs exhibit enriched integrin expression, including high expression of the laminin-binding α6-integrin (Fernandez-Nogales et al., 2022). The enriched expression of integrins in ipsilateral RGCs strongly supports the possibility that laminins may control the number of these cells by signaling through this canonical laminin receptors. Future studies will determine which receptor mediates laminin signaling in ipsilateral RGCs.

## Acknowledgements

We thank members of the Brunken, Viczian, and Zuber labs for helpful discussions and their insights. We also thank Dr. William Brunken for providing the Lamb2^-/-^ and Lamc3^-/-^ mice and reagents to get this project started, Dr. Michael E. Zuber for critical reading of this manuscript, and Francisco J. Agosto-Perez for the Python script to blind samples for analysis. This study was supported by a NIH Research Supplement to Promote Diversity for RMDL to NEI R01 EY12676 (PI William J. Brunken), Research to Prevent Blindness Career Development Award to RIMDL, Fight for Sight Grant-in-Aid to RIMDL, a NEI R01 EY034653 to RIMDL, and an Unrestricted Grant to the Department of Ophthalmology from Research to Prevent Blindness.

---

**Supplementary Figure 1.**
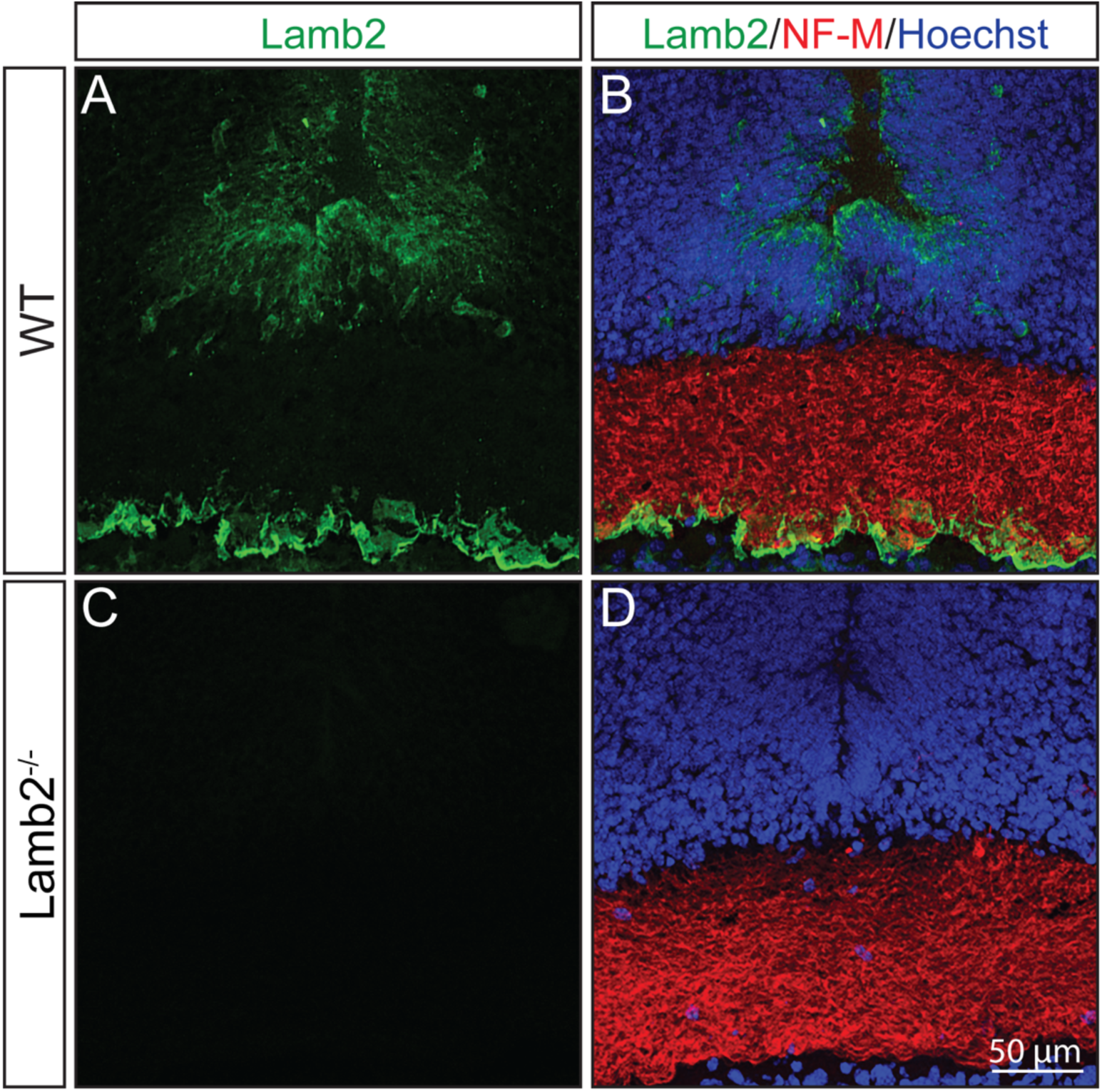
β2 laminin is not detected in Lamb2^-/-^ embryos. **(A and B)** Lamb2 protein (green) is detected in the PBM underlining the axons (red), ventricular zone, and blood vessels of the WT optic chiasm. **(C and D)** Lamb2 protein is not detected in the optic chiasm of Lamb2^-/-^ embryos. Nuclei were counterstained with Hoechst.

**Supplementary movie 1. The PBM and radial glia endfeet in the WT optic chiasm.** A 15° inclining view of the PBM and radial glia endfeet 3D reconstruction from Z-stacks of Nestin and EHS Laminin immunostained sections of the WT optic chiasm. The PBM is continuous, and the mallet-shaped radial glia endfeet connect to form the glia limitans.

**Supplementary movie 2. The PBM and radial glia endfeet in the Lamb2^-/-^ optic chiasm.** A 15° inclining view of the PBM and radial glia endfeet 3D reconstruction from Z-stacks of Nestin and EHS Laminin immunostained sections of the Lamb2^-/-^ optic chiasm. The PBM is fragmented and nearly absent. The radial glia endfeet have lost their characteristic mallet shape and are clumped into small groups.

**Supplementary Table 1.** Key Resources Table.

